# Spores and soil from six sides: interdisciplinarity and the environmental biology of anthrax (*Bacillus anthracis*)

**DOI:** 10.1101/165548

**Authors:** Colin J. Carlson, Wayne M. Getz, Kyrre L. Kausrud, Carrie A. Cizauskas, Jason K. Blackburn, Fausto A. Bustos Carrillo, Rita Colwell, W. Ryan Easterday, Holly H. Ganz, Pauline L. Kamath, Ole Andreas Økstad, Wendy C. Turner, Anne-Brit Kolstø, Nils C. Stenseth

**Affiliations:** National Socio-Environmental Synthesis Center (SESYNC), University of Maryland, Annapolis, Maryland 21401, USA; Department of Biology, Georgetown University, Washington, D.C. 20057, USA; Department of Environmental Science, Policy, and Management, University of California, Berkeley, 130 Mulford Hall, Berkeley, CA 94720, USA; School of Mathematical Sciences, University of KwaZulu-Natal, Durban, South Africa; Centre for Ecological and Evolutionary Synthesis (CEES), Department of Biosciences, University of Oslo, PO Box 1066 Blindern, N-0316 Oslo, Norway; Spatial Epidemiology & Ecology Research Lab, Department of Geography, University of Florida, Gainesville, FL, USA; Emerging Pathogens Institute, University of Florida, Gainesville, FL, USA; Department of Epidemiology & Department of Biostatistics, School of Public Health, University of California, Berkeley, 50 University Hall #7360, Berkeley, CA 94720-7360, USA; CosmosID Inc., Rockville, MD 20850, USA; Center for Bioinformatics and Computational Biology, University of Maryland Institute for Advanced Computer Studies, University of Maryland, College Park, MD 20742, U.S.A.; Bloomberg School of Public Health, Johns Hopkins University, Baltimore, MD 21205, USA; UC Davis Genome Center, University of California, Davis, 451 Health Sciences Drive, Davis, CA 95616, USA; School of Food and Agriculture, University of Maine, 5735 Hitchner Hall, Orono, ME 04469, USA; Centre for Integrative Microbial Evolution and Section for Pharmaceutical Biosciences, School of Pharmacy, University of Oslo, PO Box 1068 Blindern, N-0316 Oslo, Norway; Department of Biological Sciences, University at Albany, State University of New York, 1400 Washington Avenue, Albany, NY 12222, USA

**Keywords:** Anthrax, *Bacillus anthracis*, *Bacillus cereus*, Etosha National Park, environmental transmission, interdisciplinarity, disease ecology, eco-epidemiology

## Abstract

Environmentally Transmitted Diseases Are Comparatively Poorly Understood And Managed, And Their Ecology Is Particularly Understudied. Here We Identify Challenges Of Studying Environmental Transmission And Persistence With A Six-Sided Interdisciplinary Review Of The Biology Of Anthrax (*Bacillus Anthracis*). Anthrax Is A Zoonotic Disease Capable Of Maintaining Infectious Spore Banks In Soil For Decades (Or Even Potentially Centuries), And The Mechanisms Of Its Environmental Persistence Have Been The Topic Of Significant Research And Controversy. Where Anthrax Is Endemic, It Plays An Important Ecological Role, Shaping The Dynamics Of Entire Herbivore Communities. The Complex Eco-Epidemiology Of Anthrax, And The Mysterious Biology Of *Bacillus Anthracis* During Its Environmental Stage, Have Necessitated An Interdisciplinary Approach To Pathogen Research. Here, We Illustrate Different Disciplinary Perspectives Through Key Advances Made By Researchers Working In Etosha National Park, A Long-Term Ecological Research Site In Namibia That Has Exemplified The Complexities Of Anthrax’S Enzootic Process Over Decades Of Surveillance. In Etosha, The Role Of Scavengers And Alternate Routes (Waterborne Transmission And Flies) Has Proved Unimportant, Relative To The Long-Term Persistence Of Anthrax Spores In Soil And Their Infection Of Herbivore Hosts. Carcass Deposition Facilitates Green-Ups Of Vegetation To Attract Herbivores, Potentially Facilitated By Anthrax Spores’ Role In The Rhizosphere. The Underlying Seasonal Pattern Of Vegetation, And Herbivores’ Immune And Behavioral Responses To Anthrax Risk, Interact To Produce Regular “Anthrax Seasons” That Appear To Be A Stable Feature Of The Etosha Ecosystem. Through The Lens Of Microbiologists, Geneticists, Immunologists, Ecologists, Epidemiologists, And Clinicians, We Discuss How Anthrax Dynamics Are Shaped At The Smallest Scale By Population Genetics And Interactions Within The Bacterial Communities Up To The Broadest Scales Of Ecosystem Structure. We Illustrate The Benefits And Challenges Of This Interdisciplinary Approach To Disease Ecology, And Suggest Ways Anthrax Might Offer Insights Into The Biology Of Other Important Pathogens. *Bacillus Anthracis,* And The More Recently Emerged *Bacillus Cereus* Biovar *Anthracis*, Share Key Features With Other Environmentally-Transmitted Pathogens, Including Several Zoonoses And Panzootics Of Special Interest For Global Health And Conservation Efforts. Understanding The Dynamics Of Anthrax, And Developing Interdisciplinary Research Programs That Explore Environmental Persistence, Is A Critical Step Forward For Understanding These Emerging Threats.

## I. Introduction

The East Asian parable of the six blind sages, perhaps best known in the Western canon from John Godfrey Saxe’s “The Blind Men and the Elephant,” is an apt metaphor for the process of interdisciplinary research and multidisciplinary synthesis. In the story, the sages each attempt to describe an elephant, a new and terrifying beast they have never encountered, based solely on what they can feel. The first touches the elephant’s side and describes it like a wall; the second feels the tusks and concludes an elephant is like a spear, and so on. Saxe’s penultimate verse thus concludes, *“….each was partly in the right, / and all were in the wrong!”*

Every new epidemic and (re-)emerging pathogen represents a challenge for medical communities, and like the blind sages, each of these diseases draws researchers together to assess the nature of the beast. The sages brought to the table have of course changed over time: since the Modern Synthesis in evolutionary biology, ecologists have come into play as important counterparts of the “traditional” disease research fields, joining the ranks of microbiologists, immunologists, epidemiologists, evolutionary biologists, and clinicians (among many others). Paradigms for collaborative research like EcoHealth and One Health bring these disparate groups together to achieve interdisciplinary synthesis^1–4^—an important step towards outbreak preparedness, given that the majority of emerging pathogens have some sort of environmental origin. On a global scale, the majority of recently emerging human diseases, including those with environmental reservoirs, originate in animal populations (zoonoses) and, of those, an estimated 70% originate in wildlife.^5^ Particularly challenging to study are pathogens that blur the boundaries between direct transmission and indirect modes, including vector-borne transmission and transmission from biotic (wildlife) or abiotic (water or soil) reservoirs. In response to the significant role ecology plays in these modes, multidisciplinarity and interdisciplinarity aim to integrate ecology into disease prevention and break down the barriers that prevent meaningful communication between the “blind sages.” A number of recent high profile works have recently called for better integration of ecosystem research into disease-management efforts^6,7^, and the need to increase interdisciplinary interaction has been recognized for more than a decade^3,8^, as evidenced by a number of publications over two decades that call for removing disciplinary barriers in disease research.^1,3,4,8–13^

In the most pronounced success stories, the synthesis of interdisciplinary research has enabled two key advancements: statistical forecasting methods that enable anticipation of outbreaks, based on environmental and social data; and ecologically inspired tools for intervention to mitigate and sometimes prevent outbreaks (Figure 1). For example, cholera (*Vibrio cholerae*) was once thought to be only directly transmitted human-to-human via contaminated water (a mechanical vector), until 1983 when a team of ecologists and microbiologists discovered that copepods are aquatic hosts of cholera bacteria.^14^ Through continued collaboration among ecologists, microbiologists and clinicians, this discovery eventually enabled outbreak forecasting based on climatic data and led to the implementation of simple filtration methods that reduce case burden by as much as 48%.^15,16^ A combination of oral vaccines, water filtration techniques, improved sanitation, and predictive modeling has made the ongoing seventh global pandemic of cholera more manageable than ever before.^14–16^

**Figure 1.**
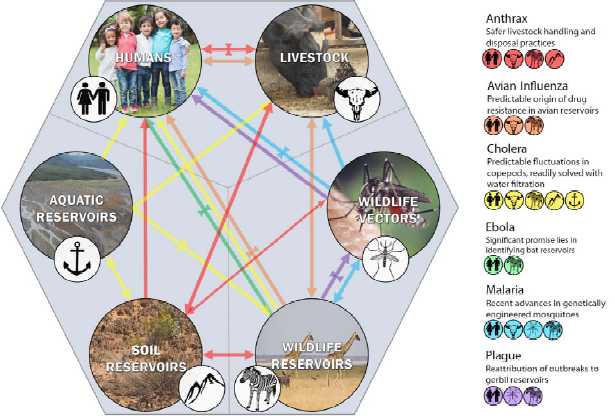
A One Health approach applied to disease systems, showing the complexity of human interactions with livestock, wildlife, and environmental reservoirs (including aquatic reservoirs, soil, and plants). Arrows represent the directionality of transmission or spillover from one compartment to another, highlighting that each disease has a unique complexity. Major discoveries presented earlier that help contain disease are shown on the transmission pathway (as bowties) they most significantly affect. Viewing diseases as organisms in their own right, navigating this web, provides a more holistic and appropriate view than only considering the human angle.

Wilcox & Colwell proposed a “cholera paradigm” for interdisciplinary research based on these advances, arguing that even for the most complex and challenging-to-predict systems, synthesis work focused on elucidating multi-component life cycles can help develop both predictive tools and prevention or control measures.^15^ But among zoonotic diseases, cholera is characterized by a simple ecology relative to its human health burden, and the discovery of its copepod host was ultimately enough to revolutionize interventions and predictions. Compared to cholera, many pathogens are still poorly understood. Newer or neglected diseases tend to show less-integrated clinical and academic knowledge; and diseases with an uncertain ecology are particularly difficult to control. Generalist pathogens with a complicated or uncertain natural history pose a particular problem for predictive work, and more often than not, the most limiting factor is a dearth of research on their ecology. In a similar undertaking to Wilcox & Colwell’s study, we demonstrate use a multidisciplinary framework to illustrate how “six blind scientists” from different disciplines would characterize recent developments in research on anthrax (B. *anthracis)*, a generalist pathogen with a more complex life cycle than cholera. We endeavor to illustrate how and why the ecology of diseases like anthrax is comparatively understudied and undersynthesized, and how interdisciplinary synthesis that includes ecology is especially important for pathogens like anthrax, precisely because there are so many unknown elements of their complex, nonlinear dynamics.

## II. Anthrax: A Case Study in Slow Integration

Anthrax is a zoonosis caused by the gram-positive bacterium, *Bacillus anthracis*, that primarily infects ungulates; other mammals, including humans, tend to be incidental hosts. Transmission takes place through several pathways, the primary one for ungulates being ingestion of *B. anthracis* spores during feeding at carcass sites.^17–19^ Other potential pathways include ingesting emesis and feces deposited by necrophagous flies on vegetation after these flies have fed on hosts that have died of anthrax^20^; inhaling anthrax spores that have become airborne, (in nature occurring from dust bathing hosts, though recent evidence cast doubt on this^21^); waterborne transmission from waterholes and temporary ponds^22^; cutaneous routes, which account for the majority of human clinical cases globally^23^; and gastrointestinal infections from eating infected meat and blood directly.^17,18,24,25^ In some regions, anthrax outbreaks are a consistent and predictable feature of ecosystems and occur regularly on a seasonal cycle; in other settings, epizootics are infrequent or rare events, and can be responsible for mass die-offs among wildlife and livestock.^17,26,27^

While epidemic and pandemic threats like Ebola and Zika rightly attract substantial attention in both disease ecology and the public health realms, anthrax maintains a lower profile in terms of cumulative impact on global health, normally only making headlines in the context of bioterror scares or rare mass die-offs of wildlife. Despite its global presence, and its wide range of suitable hosts, little is known about the dynamics of anthrax in most of the ecosystems and hosts it infects. The pathogen is best studied in a handful of regions, in particular the Midwestern United States, the former Soviet Union, and sub-Saharan Africa. The insights we review here come from two decades of work based especially in Namibia’s Etosha National Park, a savannah ecosystem that is host to high ungulate diversity and endemic anthrax. Data collection regarding anthrax outbreaks in bovids, elephants, zebra, and other mammals in Etosha began in the 1960s, and has provided one of the most continuous sources of documented anthrax dynamics in any natural system. Interdisciplinary work has emerged from treating Etosha as a window into the complex and often hidden dynamics of anthrax spores in the environment, and to date represents one of the most successful ventures to better understand the ecology of the disease. Here, we review six disciplinary perspectives (Figure 2) on anthrax dynamics in sub-Saharan Africa, especially in Etosha^28^; and highlight how each has contributed to scientific understanding about anthrax’s life cycle.

**Figure 2.**
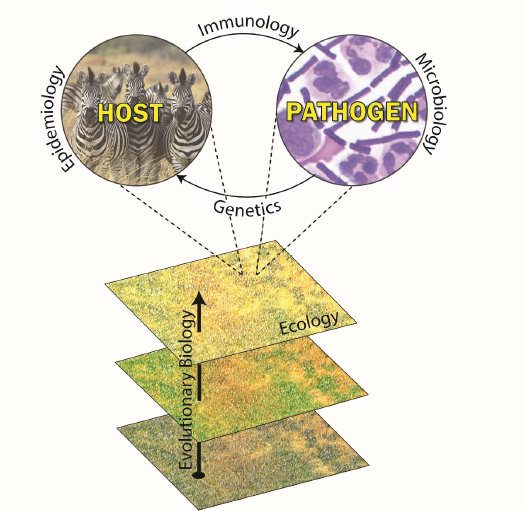
Different disciplinary perspectives fit together to provide a holistic perspective on pathogen ecology. Some occur at multiple scales (e.g. while we aggregate genetics, genomics, and evolution in the main text, their study may occur somewhat separately at different scales). The sixth presented in our main text, clinical and public health, occurs in parallel with all the processes depicted.

### (1) Microbiology

Louis Pasteur first proposed that carcass sites could function as the main route of anthrax transmission^29^. Long considered an obligate pathogen, *B. anthracis* was thought to replicate only within a vertebrate host, where conditions were conducive for proliferation of vegetative cells.

When the host succumbs to the anthrax infection, vegetative cells of *B. anthracis* are released (along with blood and other body fluids) into the environment, and produce infectious spores capable of long-term survival. The environmental maintenance of anthrax is the least understood part of its life cycle; despite the central role of the environment in its transmission, surprisingly little is known about the survival and activity of *B. anthracis* outside of a host. Three fundamental questions have been at the heart of most research on this topic: where and for how long can *B. anthracis* persist in the environment, is it capable of germinating in the soil under any normal conditions, and how does it interact with other microorganisms and plants?

How long do anthrax spores persist in the environment? Evidence from a cross-scale study by Turner & Kausrud *et al.^19^* suggests that cultivable *B. anthracis* spore concentrations exponentially decline over time in the soil. In the first two years after a carcass site is formed and spores are deposited, any grazing at carcass sites is likely to infect hosts, even just from eating above-ground plant parts of grasses; anthrax spores tend to decay to negligible concentrations on grasses after more than two years. However, after 2-4 years, anthrax spores still persist in the soil at high enough concentrations that herbivores ingesting soil directly, or indirectly along with grass roots, are probably still at significant risk of infection. Overall, transmission through grazing appears to be most likely in the 1-2 year window when grass growing at former carcass sites is more abundant and nutritious.^19^ Nevertheless, data from Etosha shows that spores can be detected for more than seven years after decomposition of the carcass^22^, and it is likely that longer-term persistence drives anthrax dynamics in other more episodic systems (like western Canada’s Wood Bison National Park), where exponential rates of spore decay in cooler, more heavily vegetation-covered, less intensively radiated soils may be substantially slower than in Etosha. In some natural conditions, spores are known to persist for decades; spores of the Vollum strain of *B. anthracis* were detected more than forty years after soils were experimentally inoculated at Gruinard Island.^30^ Similarly, re-emergence of anthrax in reindeer in the Yamal region of Siberia in 2016 more than seventy years after the last known case, together with sporadic cases originating from unknown environmental sources in Sweden, strongly suggests that persistence times can exceed one or more centuries under certain conditions. The rate at which a spore pool decays likely depends heavily on local environmental conditions: for example, larger concentrations of spores are found in soils having slightly alkaline pH, higher organic matter and higher calcium content.^18^ Features of the exosporium also have been shown to affect the ability of *B. anthracis* spores to bind to different soil types.^31^

Originally, data from spore-contaminated soil samples in some areas indicated that *B. anthracis* has a tendency to lose the pXO2 plasmid (encoding the poly-γ-D-glutamic acid capsule) over time (5-8 years), suggesting that at least a minimum amount of replication^32^, and therefore genetic evolution, takes place in the soil environment^33^; subsequent work has confirmed that *B. anthracis* can in fact replicate in the soil. However, the conditions under which this can happen are highly controversial. Van Ness proposed that under conditions of alkaline pH, high soil moisture, and the presence of organic matter, *B. anthracis* can maintain a high population density by replicating in the environment^34^. However, this “incubator hypothesis” remains controversial because Van Ness did not provide empirical support. Moreover, laboratory studies suggest that although vegetative cells may potentially flourish outside of a host, their survival in the environment may be significantly influenced by antagonistic interactions with other microbes. In early experimental studies, Minett and Dhanda^35^ and Minett^36^ found that *B. anthracis* spores germinate and multiply in moist sterile soil but not in soil with a microbially-diverse population, and suggested that bacterial antagonists may restrict its activity. In further work, several species of soil bacteria were found to impede growth of vegetative *B. anthracis* cells^37^, and a separate study found that some bacteria typically present in soil inhibited multiplication of vegetative *B. anthracis* cells in unsterilized soil.^38^ (For comparison, the closely related *B. thuringiensis* also persists in the soil for extended periods^39^, and may be capable of germinating in the soil, but is similarly often outcompeted by other bacterial species *in situ*, and grows better in sterilized soil.^40,41^) Conversely, the effect of *B. anthracis* on other soil bacteria has also proved interesting, if complex. In a groundbreaking study, Saile and Koehler demonstrated that *B. anthracis* germinates in the rhizosphere of plants (but did not find evidence for multiplication), suggesting replicative cycles in the rhizospere of grass plants to play a potential role in the evolution of *B. anthracis*, as it does for other members of the *B. cereus* group^42^, including *Bacillus cereus*^43^ and likely *Bacillus thuringiensis.*^44–46^ In this respect, it is interesting to note that *B. subtilis* recently has been shown to protect plants against bacterial pathogens in the plant rhizosphere, and that the protective effect requires biofilm formation.^47,48^ Although knowledge is lacking about a potential role for *B. anthracis* biofilm formation, in the rhizosphere or during infection, the bacterium is capable of biofilm formation *in vitro.^49^*

Recent ecological studies have shown that *B. anthracis* also interacts more broadly with some other members of the grassland-soil community, including plants^42^, earthworms^29,50,51^, and soil amoeba.^52^ Pasteur was the first to propose that earthworms vector *B. anthracis* from buried livestock carcasses, and he isolated *B. anthracis* from earthworms collected in surface soils at a burial site.^29,50^ Following up on these observations, Schuch and Fischetti^51^ found that bacteriophages can generate phenotypic changes in *B. anthracis* that enable it to persist as an endosymbiont in earthworms, and to act as a saprophyte in soil and water. Under simulated environmental conditions, Dey *et al.^52^* showed that a fully virulent Ames strain (pXO1+, pXO2+) of *B. anthracis* germinates and multiplies intracellularly within a free-living soil amoeba living in moist soils and stagnant water, and that the pXO1 plasmid was essential for growth. This may indicate that amoebae and possibly other soil-borne protozoa contribute to *B. anthracis* amplification and spore persistence by providing an intracellular environment that allows the completion of a life cycle from spore germination through vegetative growth and back to spore formation. Several other mammalian pathogens (e.g. *Francisella* spp. and *Legionella pneumophila*) are known to be capable of parasitic or symbiotic relationships with amoeba^53,54^, and there may be a direct link between a pathogen’s ability to survive within amoeba and its ability to survive encounters with primary phagocytic cells (macrophages and neutrophils). Some researchers have even gone as far as to suggest that co-evolution between these bacteria and amoeba seems to have allowed the parasitism of mammals.^54^

### (2) Genetics, Genomics, & Evolution

Bacillus anthracis shares a common chromosomal framework with all main species of the *Bacillus cereus (sensu lato*) super-species group, including the opportunistic soil bacterium *B. cereus*, and the entomopathogenic *B. thuringiensis*, which (because of analogous plasmids occurring throughout the group) sometimes leads to blurred species boundaries.^55–60^ The chromosomal elements principally separating classic *B. anthracis (senso stricto*) from the other closely related species are: 1) the presence of four distinctive chromosomal prophage elements; 2) a specific, inactivating nonsense mutation in the transcription factor PlcR, a positive regulator mainly of chromosomally encoded extracellular virulence factors that are important during mammalian and insect infections by *B. cereus* and *B, thuringiensis*; and 3) being part of the genetically monomorphic *B. anthracis* cluster by phylogenetic analysis.^55^ In addition, *B. anthracis* requires two large plasmids for full virulence: pXO1, which encodes the anthrax toxins, and pXO2, which encodes the protective poly-γ-D-glutamate capsule. Large-scale, whole-genome sequencing studies suggest there has been no recent large-scale gene loss in *B. anthracis* or unusual accumulation of non-synonymous DNA substitutions in the chromosome.^57^ The fact that *B. anthracis* spends large parts of its evolutionary time as a dormant spore (on average the bacterium carries out 0.28-1 generation per year^61^), presumably contributes to its highly monomorphic nature. During infection of a host, however, mutations accumulate, and it is thought that genetic evolution of *B. anthracis* is mainly limited to the roughly week-long periods between exposure and host death, estimated to cover 20-40 bacterial generations.^62^ Selection in *B. anthracis* can subsequently act upon phenotype with regards to the spore, based on mutations acquired during its last infective cycle.^18^

Furthermore, based on genetic data, the *B. anthracis* species can be divided into three distinct subpopulations, the A-, B-, and C-branch. C-branch isolates seem to be strikingly rare, and B-branch isolates are geographically restricted to southeastern Africa and western Europe, while A-branch isolates have been hugely successful with respect to lineage multiplication and geographical spread worldwide.^63^ In addition, a number of different genotypes of *B. anthracis* may be present in endemic regions^64^, which may also potentially give rise to co-infection with more than one genotype.^65^ In the context of the relationship between *B. anthracis* genetics and transmission, however, little is known regarding the relationship between different strains (genotypes) and virulence levels, and hence estimates of LD_50_ by and large fail to account for variation between different host species, or between different immunological and physiological states of individuals.

Although *B. anthracis* has been previously suggested to have undergone a genetic bottleneck fixating it as a genetically monomorphic pathogen^66^, it is interesting to note that consolidation of clinical and eco-evolutionary (DNA sequencing) data during the past decade indicates that what presents as “anthrax” also may include specific isolates of *B. cereus*, causing opportunistic anthrax-like infections in humans and great apes.^67–70^ These *B. cereus* group variants, such as *B. cereus* G9241 and *B. cereus* biovar *anthracis* (including the first identified CI and CA strains) ^67,69,70^, have been isolated from cases of human (USA) and animal (West Africa) infections, respectively, involving anthrax-like symptoms and disease. These cases are genetically different from classical *B. anthracis* in that they carry variants of the pXO1 and pXO2 (or another capsule-producing) plasmids in a *B. cereus* chromosomal background, and clearly constitute evolutionary distinct lineages in the *B. cereus* group phylogenetic tree representing different evolutionary origins from classical *B. anthracis.^55^* Specifically, the G9241 strain encodes a non-mutated and potentially functional PlcR regulator in the chromosome, thereby imposing a *B. cereus* phenotype (hemolytic, motile).^67^ Conversely, *B. cereus* biovar *anthracis* strains carry a *plcR* mutation in a location closer to the 3' end of the gene than in classical *B. anthracis*, but resulting in an essentially *B. anthracis* phenotype (non-hemolytic, non-motile). Both *B. cereus* G9241 and biovar *anthracis* strains are capable of producing a second type of capsule (hyaluronic acid) from a locus (*hasACB*) residing in pXO1^71^, which in classical *B. anthracis* carries an inactivating single nucleotide deletion.^72^

The recurring presence of such strains in natural settings shows that active transfer of pXO1 and pXO2 variants was not a one-off evolutionary event for a lineage leading up to *B. anthracis.* On the contrary, these observations allow for the possibility of pXO1 and pXO2 transfer into *B. cereus* having taken place as multiple independent evolutionary events, leading to multiple lineages of strains with the capacity to evolve into contemporary strains of variable genetic composition, but producing similar anthrax-like symptoms during human or animal infections. However, it is interesting to note that the *B. cereus* biovar *anthracis* isolates from animals in Cote d’Ivoire, Cameroon, Central African Republic and Democratic Republic of Congo, similar to *B. cereus* G9241, appear to belong to the same main phylogenetic clade (B. *cereus* group clade I^55,59^), and that the pXO1 and pXO2 plasmids hosted by the biovar *anthracis* strains followed the chromosomal phylogenies with the same branching order.^28^ These data indicate that both the chromosomes and the plasmids in the isolated strains are closely related. In addition, strains isolated from Cote d’Ivoire and Cameroon were tested in toxicity and vaccination experiments and found to be just as toxic as fully virulent *B. anthracis*, and protective vaccination in mice and guinea pigs were just as efficient as for *B. anthracis*.^71^ Recent epidemiological studies further suggest that these strains are more widespread than initially thought, constituting major drivers of wildlife mortality in the African rainforest rather than representing isolated, single cases of wildlife disease.^26^

### (3) Immunology

Anthrax infections end either in recovery, or death of the host. When *B. anthracis* spores are ingested, the spores germinate into fast-multiplying vegetative forms that produce three soluble factors that assemble to form toxic complexes: edema factor (EF), an adenylate cyclase that impairs immune cell function^73–76^; lethal factor (LF), a zinc metalloprotease that cleaves MAP-kinase-kinases thereby suppressing production of several types of cytokines and immune cell functions^77–82^; and protective antigen (PA), which complexes with the other two factors and allows them to enter host cells through oligomeric PA pores.^83^ The PA and LF bind to form the anthrax lethal toxin (LeTx), the key virulence factor of *B. anthracis* that kills macrophages and dendritic cells through a caspase-1-dependent cell death program known as pyroptosis.^84^ The collective actions of these toxins may ultimately result in the peracute-to-acute death of susceptible hosts from edema, vascular collapse, and inflammation, combined with an overwhelming septicemia of up to 10^9^ bacterial cells per milliliter of blood.^85–92^

Lethal dose is a host-population concept, and within populations, hosts will vary in their susceptibility to anthrax because of inherited genetic factors, as well as current immunological status, coinfection, and physiological condition. Most studies regarding host immune responses to anthrax have been conducted in laboratory settings. These studies have demonstrated that humoral immunity, particularly against the PA toxin, plays a very important role in a host’s fight against anthrax; the presence of anti-PA antibodies appears to be essential for adaptive protection, and several studies have demonstrated that the magnitude of a host’s anti-PA IgG antibody titer is correlated with level of protection against the disease.^85–92^ Furthermore, anthrax vaccine studies have indicated that T cells may also play a role in immunity to anthrax.^93^ While anthrax spores require phagocytosis by macrophages for germination, macrophages have also been found to play a primary role in limiting and clearing anthrax infection.^94,95^ Following infection, macrophages engulf and destroy invading pathogens, recognizing *B. anthracis* through toll-like receptor 2 (*TLR2)^96^* Genetic studies in mouse models and cell lines have identified several host genes that modulate susceptibility to *B. anthracis* infection and support a multigenic contribution to the host response.^97^ The myeloid differentiation factor (*myD88)*, a downstream mediator of the TLR pathway, has been shown to confer susceptibility to anthrax^98^, and polymorphisms in the *Nlrp1b (Nalp1b*) gene have also been shown to influence susceptibility to the anthrax toxin in mouse macrophages^99^ and human fibroblasts.^100^ However, other studies have demonstrated that the LT-sensitive *Nlrp1b* allele induces early inflammation that protects against anthrax.^101,102^ *TEM8* and *CMG2* genes encode host transmembrane proteins that function as anthrax LeTx receptors^103,104^, binding with PA and mediating delivery of LF into host cells. *CMG2* has a considerably higher affinity for LeTx than does *TEM8*, and CMG2-null mice are highly resistant to *B. anthracis* infection.^105^ In fact, *CMG2* variation significantly alters toxin uptake and sensitivity in humans, with lethality differing up to 30,000-fold among cells from people of different ethnic backgrounds.^106^

While these studies provide a wealth of mechanistic knowledge about the host’s immunological response to anthrax, the scaled application of that information to anthrax dynamics is far more complex—especially as laboratory studies on mice can only reveal so much about the dynamics of infection in large herbivores. To remedy this, field studies are needed to address gaps in our understanding of anthrax infections, and of how immunology scales up to produce broader eco-epidemiological patterns. One particularly important problem is the effect that sub-lethal doses may have in promoting adaptive immune responses to anthrax.^107–111^ Recent evidence suggests that sublethal anthrax infections in species known for high apparent mortality—including the herbivores like zebra and springbok that are most abundant and most important in anthrax outbreaks in Etosha—are more common than previously thought.^112^ In fact, frequent anthrax contact can act as an immunity booster in both carnivores and herbivores, strengthening their anti-anthrax protection over time and possibly lessening the overall morbidity and mortality within the population (endemic stability).^111–113^ In contrast, evidence suggests sub-lethal infections of *Bacillus cereus* biovar *anthracis* in primates are incredibly rare^114^, which has dramatic implications for the long-term survival of chimpanzee populations in West Africa.^26^

There are likely to be important causal relationships between the mammal species present in a community and the dynamics of anthrax (and perhaps vice versa?) through their interspecific immunological variation, but not enough is known about most mammal species to hypothesize cross-system rules at the present time.

A similar key problem is that anthrax infections exist in an ecosystem of pathogens, and the within-host role that co-infection plays in anthrax immunology is complicated at best. Field studies in Etosha have shown apparent tradeoffs in zebra between Th-1 and Th-2 type immune responses, where Th-2 responses seem to peak in the wet season in response to gastrointestinal helminths.^115^ These immune responses appear to make zebra and springbok more tolerant of helminth infections when they peak during the wet season, decreasing host resistance to anthrax infection and thereby potentially contributing to the overall seasonality of anthrax outbreaks.^116^ Geographic variation in local pathogen communities might therefore influence mortality and outbreak frequency of anthrax across systems, and while this kind of community ecology approach is sorely needed in many aspects of disease ecology^6^, it is again hard to imagine a future with sufficient data availability to convincingly address these questions at broad scales.

### (4) Ecology

Landscape ecology research on anthrax has had the greatest successes by studying the locations and processes of herbivorous hosts that have died of anthrax (Figure 3). These carcass sites act as “locally infectious zones” (LIZs)^117^, and come to have a demography of their own as these zones appear and fade over time. Rather than passively acting as a fomite, evidence suggests that anthrax carcass sites have a complex set of biotic interactions that determine their persistence and infectiousness throughout a landscape. Nutrient deposition from carcass decomposition appears to be the primary correlate of overall plant growth in green-ups; zebra carcasses in Etosha substantially increase soil phosphorus and nitrogen that persists over at least three years.^19^ However, experimental evidence suggests *B. anthracis* spores also directly facilitate the germination of grass seeds.^118^ The mechanism through which that occurs is still uncertain (and requires further investigation), but Saile and Koehler demonstrated that anthrax germinates in the plant rhizosphere^42^; as *B. anthracis* is a member of the *B. cereus* group, it is possible that *B. anthracis* retains some of its ancestral capabilities to engage in beneficial plant-microbe interactions. There has been no evidence, however, that plants facilitate persistence of *B. anthracis* in the soil, potentially suggesting that its association with vegetation may serve to attract herbivores that ultimately become infected and spread the pathogen across the landscape.^118^

**Figure 3.**
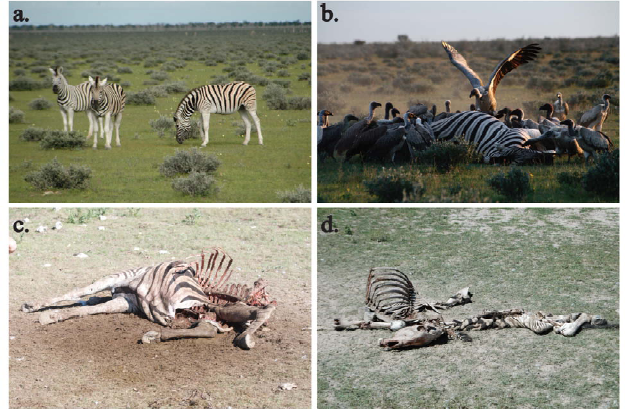
The life cycle of anthrax in Etosha, viewed from the perspective of zebra (a), the most common host. Zebra become infected while grazing, dying within approximately a week and immediately attracting scavengers (b) that quickly open a carcass, depositing spores into the ground. During the early stages of a carcass site, herbivores can fairly easily identify and avoid partially decomposed carcasses (c), but as carcasses slowly blend into the environment over a period of years and vegetation returns (d), herbivores return and once again become infected.

Herbivores in Etosha face a tradeoff between the benefits of foraging at green-ups and the obvious costs associated with lethality, with possible selective pressure acting on foraging strategy.^19,119^ Evidence based on camera trap data from Etosha suggests that most herbivores avoid carcass sites early in the first year of establishment because they are denuded of vegetation by scavengers, but that they in fact favor green-ups following the first rains after the nutrition influx from the carcass. This likely contributes to the importance of the 1-3 year window after establishment in transmission.^19^ Anthrax dynamics are also seasonal within years, peaking in March-April, which more generally aligns with the later part of the warm wet season in Etosha. While some research previously suggested that nutritional stress might drive the seasonality of anthrax, evidence directly contradicts the idea that nutritional stress is worse during the anthrax season.^116,120^ Instead, it appears that soil ingestion increases during the wet season for a handful of species including zebra, directly increasing anthrax exposure; in contrast, elephant deaths (while rare) in fact peak in October-November, suggesting there are interspecific heterogeneities in exposure pathways that still require investigation.

Similarly, interspecific variation in movement still requires investigation, given the wealth of movement data collected in Etosha over the past two decades^121–126^; for example, elephants largely migrate away from the known anthrax areas of Etosha during the anthrax season, and return in the dry season. Intraspecific variation also requires further investigation study; evidence suggests that there may be a link between partial migration of zebra herds in Etosha and avoidance of the anthrax season. It has been suggested that this phenomenon could be linked to dominance structure, as dominant groups migrate, while resident (submissive) herds are encouraged to stay by decreased competition.^123^ Movement data from Etosha has already been used to help develop anthrax-relevant analytic tools^122^ and simulations^121^, and while some work in other systems has used movement data and tools to help map the link between environmental suitability and host exposure^127^, similar work is still needed in Etosha (as it is needed in any system with the potential for transmission at the wildlife-livestock interface, where these tools are often the most useful for answering applied questions^128^).

How do non-herbivorous mammals affect anthrax dynamics? Some work had suggested that scavengers might play an important role in the dispersal and creation of LIZs, but work in Etosha has often proved counter to those ideas. Previous theory had suggested vultures might disperse bacteria from carcass sites to their nesting sites and thereby help spread disease, based on anecdotal evidence^129^, examination of faeces and contaminated water ^17,27^, and experimental work on anthrax spore passage through vulture digestive tracts.^130,131^ However, research in Etosha failed to find higher *B. anthracis* concentrations at vulture nests, perhaps because vultures’ acidic droppings produce soil unsuitable for anthrax spores.^132^ Furthermore, prior work indicated that during the first 72 hours after carcass deposition, if a carcass remains unopened, vegetative cells fail to sporulate, ending the life cycle. ^17,27,133^Consequently, scavengers seemed likely to play a significant role in anthrax dynamics, by tearing open carcasses and promoting blood flow into the soil.^18^ In contrast, an alternative hypothesis suggested that scavengers— especially vultures and other birds, which are less prone to anthrax-related deaths due to acquired immunity^108^—could “cleanse” carcass sites, reducing LIZ formation and establishment. However, experimental exclosure of scavengers from zebra carcasses in Etosha revealed that scavengers had no effect on soil spore density, failing to find evidence for either hypothesis, and further challenging the role of scavengers in anthrax ecology.^134^ In other systems, with a different assemblage of scavenger species, the role of scavengers in spore dispersal or LIZ formation may be different than in Etosha, and therefore could require further investigation.

### (5) Epidemiology

The epidemiology of directly-transmitted pathogens draws upon ecology, particularly behavioral ecology, to better understand how susceptible and infected individuals come into contact with one another; by comparison, the epidemiology of environmentally transmitted pathogens, such as anthrax, requires a much wider understanding of the relevant host and pathogen ecologies, particularly the interactions of hosts and pathogens within their environments.^117^ Thus, in the case of indirect transmission, it is more difficult to separate the ecological and epidemiological components. Appropriately complex epidemiological models^135^ are needed that can unpack different aspects of transmission, including the dose of pathogen that hosts are exposed to, the immunological variation between individual hosts and between species, and the interplay between the two.^136,137^ Modeling the dose-exposure process requires an understanding of individual pathogen shedding into the environment, the movement of susceptible individuals through space, the internal milieu of the host, and in some cases the behavior of susceptibles once the source has been encountered.^128^ However, studies that explicitly consider the movement of individuals across landscapes, the ability of pathogens to persist in the environment, the immunological status of susceptible individuals, and issues of dose or prior low-dose exposure are rare.

Studies in Etosha have provided the tools to begin to develop models that cross these different scales, and thereby unravel the false “lethal dose paradox,” in which the experimentally determined lethal dose required to kill herbivores appears to be far higher than would be encountered in nature.^22^ Work that combines field experiments and modeling shows that, especially in the first two years after deposition, carcasses should provide ample infection risk for grazing herbivores, even when soil ingestion is minimal.^22^ Even though spore concentrations begin to decline rapidly after two years, they may still be sufficient to produce sporadic outbreaks, especially in drought years that intensify herbivore soil contact during grazing. These small outbreaks may set off (and often precede) epidemic years, illustrating how the long tail of
LIZ persistence could ultimately play an important role in long-term anthrax dynamics^22^. Studies like this allow future work that considers the epidemiology of anthrax at large scales more explicitly, such as by using agent-based models to simulate the effects of host heterogeneity and landscape structure on outbreak dynamics, and measuring the relative utility of different host movement metrics as predictors of anthrax risk.

While the epidemiology of anthrax in Etosha is better understood in light of recent modeling studies^22,117^, Etosha is only a single landscape, and anthrax outbreaks behave very differently around the world. The reasons for differences in the frequency, timing, and intensity of anthrax outbreaks globally are poorly understood, but stem from some combination of microbiological, immunological, and ecological factors discussed above.^117^ Not all possible transmission modes are important in Etosha; for example, vector enhancement by necrophagous flies has been implicated as an important mode of spores being spread from carcasses onto above-ground vegetation, but appears to play a minimal role in Etosha. However, some universal patterns can be noted. For instance, soil type and alkalinity is known to affect spore persistence^18^, and other climactic conditions may play roles in determining when and how often outbreaks occur. Anthrax outbreaks in the middle latitudes appear to be seasonal across host systems.^17,138,139^For example, deer outbreaks in Texas appear in summer months^17^, with the severity of outbreaks increasing in response to early and intense spring green-up.^140^ Similarly, anthrax outbreaks tend to be observed in Etosha (and elsewhere) with dry conditions following periods of intense rainfall, for a number of potential reasons, including changing animal movement patterns (without water-restrictions, animals range more widely at lower densities, whereas in drier periods, they aggregate at waterholes), changing vegetation growth or processes changing spore density on vegetation (such as splashing of spore-laden soil onto grasses)^120^, and increasing exposure to other potentially interacting microparasites and macroparasites (altering host susceptibility^116^).

While the subtleties of community ecology and herbivore movement make mechanistic modeling a challenge at the landscape level, the microbiological factors that determine spore production, dispersal, persistence and amplification will ultimately determine the location and persistence of LIZs at much more flexible spatial scales, and these abiotic constraints ultimately determine broader-scale patterns of presence or absence. Through the use of ecological niche models (ENMs), anthrax occurrence data can be used to study and predict the environmental covariates driving persistence at continental, national, and sub-national scales. By combining predictive understanding of environmental persistence (with model selection and variable selection heavily informed by studies and understanding at the microbiology and landscape ecology scale) with studies at the human-wildlife-livestock interface, regional surveillance tools can be developed that appropriately map anthrax risk.

It is worth noting that while anthrax is effectively cosmopolitan on a global scale, no global map of its distribution has ever been constructed; instead, most studies have mapped its distribution using ENMs constructed at the regional or national scale, often in close partnership with public health efforts. Studies using ENMs to map anthrax have been done in at least 12 countries, including Australia^141^; Cameroon, Chad, and Nigeria^142^; China^143^; Ghana^144^; Italy and Kazakhstan^145^; Kyrgyzstan^146^; Mexico and the United States^147^; and Zimbabwe.^148^ In many of these regions, ENMs are the most accessible statistical tool for mapping risk, and therefore guide spatial prioritization for wildlife surveillance and livestock vaccination campaigns; however, follow-up studies evaluating the efficacy of ENM-based campaigns are currently fairly lacking (a problem not unique to anthrax work). ENM-based methodology is also incredibly flexible, and can be used in combination with other tools such as resource selection functions to improve predictions of how herbivore movement drives cases, or hotspot analyses to study clusters of human and livestock case data.s^127,128^ These studies can even be used to project future scenarios, including the role climate change will play in altering anthrax transmission.^149,150^

The greatest strength of ENMs as an approach is the ability to identify and map risk patterns, based on incomplete datasets that aggregate data across species and over longer stretches of time, without an underlying mechanistic or compartmental model. For some pathogens, more mechanistic spatiotemporal models that project spillover risk may be desired, and may be more readily constructed over time as the zoonotic process becomes clearer^151–153^; but for a generalist pathogen like anthrax, the complexity of herbivore communities and of the wildlife-livestock-human interface might make these types of models impossible to develop. However, ENMs operate on a broader temporal scale than may be helpful for short-term outbreak anticipation efforts. Early warning systems (EWSs), based on leading indicators^154,155^ or on climatic covariates^156,157^, address the problem of outbreak prediction on a shorter timescale, and are a high priority for development for many diseases—but to the extent of our knowledge, these models have yet to be developed and deployed for anthrax. The existence of long-term outbreak datasets from long term research sites like Etosha or Kruger could likely support the development of these tools, but the transferability of these models to other regions would be limited both by the selection of statistical fitting procedures, and by differences in the underlying eco-epidemiological process across systems.

### (6) Public Health

Like all environmentally transmitted diseases, anthrax poses a substantial challenge for surveillance efforts, as the majority of anthrax dynamics are unobserved either due to the difficulties of studying anthrax in the soil, or the limited resources available for epizootic surveillance. The acuteness of anthrax infections makes the window of detectability very narrow for infected animals, and even once carcasses are deposited, many are not found for days or are never found at all, depending on scavenger presence and the location of death; one modeling study estimated anthrax carcass detection rates in Etosha, a well surveilled site, at roughly 25%.^158^ Even still, the Etosha site has been nearly unique in the depth and detail of coverage, with over 50 years of data collection. Anthrax is globally cosmopolitan, and outbreak data for human, wildlife and agricultural cases are most commonly collected from passive surveillance following both wildlife and livestock mortality. While these data are limited, they can be used in combination with our understanding of anthrax dynamics at local scales to create public health-relevant predictive infrastructure (especially statistical tools like ENMs that describe spatiotemporal patterns of risk), and therefore help guide the targeting of intervention campaigns.

Anthrax eradication is, for any given landscape, an essentially impossible task given the soil spore banks and the often-cryptic enzootic process. However, livestock vaccination programs, carcass removal, and avoidance of high-risk locations have been shown to greatly improve regional outcomes. Where combined with surveillance, public health and veterinary infrastructure to deal with outbreaks when they occur, regional patterns of emergence have been kept intermittent and low-impact. An anthrax vaccine is currently available for both animals and humans^89^, but its memory response—as well as the memory response to natural, sub-lethal anthrax infection—tends to remain elevated for only a few months^92,112^, the reasons for which are unknown. In addition to vaccination, control efforts focus on sanitary carcass disposal. However, in areas where dead animals will not be discovered before body fluids have leaked into the ground, the success is limited, as soil sterilization is costly and inefficient.

Around the world, control efforts are highly variable, and necessarily correspond to the local ecology. In the Etosha, control efforts centered on sanitizing of carcasses, but the remoteness of the area, lack of local firewood sources and fragility of vegetation to heavy machinery caused control efforts to be discontinued in the 1980s. Since then, anthrax has been endemic and seen as a natural part of the ecosystem with annual outbreaks in wildlife in the area. All host species seem capable of keeping stable populations however, though elephants may be vulnerable due to fluctuations in their smaller population when combined with poaching pressure. More generally in Africa, as anthrax is widely endemic, livestock vaccination is the primary method of protection from spillover of wildlife epizootics; but coverage is likely low in many populations of rural poor livestock keepers.^144^ Consumption of meat from infected livestock may be common in these populations (as it is worldwide, as a primary route of infection), especially if medical knowledge is limited in these communities; human fatalities that result may be severely underreported.^109,110,144,159^

In Russia, anthrax is known as the “Siberian plague” (сибирская язва) due to its historical high prevalence in Siberia. ^160^ It has been largely controlled in the last half century due to large-scale vaccination of domestic reindeer herds, combined with efforts at tracking and avoiding burials of infected animals. In the Yamalo-Nenets region, reindeer vaccination efforts started in 1928^160^, but were discontinued in 2007 because no new cases had been observed since 1941.^161,162^ Following unusual permafrost thawing in the summer of 2016, three simultaneous anthrax outbreaks killed almost 2500 reindeer and caused the culling of several hundred thousand more during control efforts^163^; a hundred people were hospitalized and one boy died.^164^ Even if every carcass were sanitized, new infections are likely to occur for the foreseeable future, as the spore pools of Siberia are likely to persist for decades more (making some land unusable for traditional reindeer herders^162^).

In the industrialized countries of western Europe, large-scale anthrax outbreaks have been absent in modern times due to sanitation and vaccination^165^, but even in Sweden the summer of 2016 saw outbreaks in domestic cattle from old environmental sources of unknown locations.^166^ The outbreaks were rapidly controlled through the normal efforts of carcass disposal, culling, and vaccinations, but it remains clear that environmental spore banks will continue to exist indefinitely even in modern agricultural areas, ready to emerge as epidemics should veterinary infrastructure falter. Similarly, in the United States, widespread access to affordable livestock vaccination since the 1950s has substantially reduce anthrax risk, but vaccination is not mandatory, and typically used in reaction to outbreaks rather than to prevent them—allowing for enough cases to maintain the pathogen in the long term. Furthermore, in regions like western Texas where anthrax is maintained by epizootics in white-tailed deer, the absence of an oral vaccine and therefore inability to vaccinate livestock substantially reduces managers’ ability to control the disease.^167^

### (7) Discussion

Data on anthrax is still incomplete, and established knowledge is subject to change in many disciplinary perspectives (Figure 4)—and consequently, the overall integration of knowledge is comparatively limited. Some basic topics still remain controversial, like the role *B. anthracis* plays in the rhizosphere or the importance of flies and scavengers across ecosystems, and will likely continue to be ongoing topics of research and debate. But our understanding of the biology of anthrax in Etosha has also significantly deepened since pioneering work in the 1970s^168^, and we contend that work from Etosha highlights the strength of interdisciplinary research on the environmental biology of neglected diseases like anthrax. The growing understanding of anthrax ecology helps explain the role it plays in the savannah ecosystem, and has helped develop better One Health type collaborative surveillance projects^169^; and the ecological and epidemiological insights we discuss above have the potential to inform tentative early warning systems, or encourage the development of ecologically-minded interventions, in similar situations.

**Figure 4.**
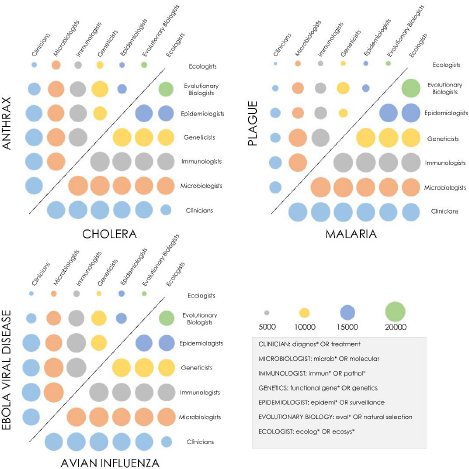
The state of interdisciplinarity in disease research, as shown by Google Scholar results for papers published 2000-2015 at the nexus of seven disciplines (an approximate method for a top-down view of literature). For some diseases, like cholera, a strong interdisciplinary focus allows ecologists and clinicians to interact at the same intensity as researchers in more closely related fields. But for other neglected diseases, like anthrax, intra-host research (microbiology and immunology especially) dominate clinical collaboration. In the poorly-integrated literature on these diseases, ecological insights translate into human health solutions in a limited way.

More broadly, work in Etosha still suggests basic rules for system dynamics, addresses basic information about pathogen biology and genetics, and offers a template for experimental and modeling work needed to understand the potentially different and unexpected eco-epidemiological characteristics of anthrax in a novel ecosystem. Even in more episodic anthrax systems, which may follow altogether different rules, research from Etosha points to the importance of understanding the factors limiting spore persistence in the soil; the relationship between anthrax and local plant communities; and the feedbacks among anthrax dynamics, host community ecology, immunity, behavior, and other local pathogens (including both related *Bacillus* and macroparasites). Expanding our understanding of variability in anthrax ecology, and plasticity (and evolutionary change) in anthrax’s life history, will help contextualize ongoing work relating its global distribution to patterns of diversification.

Developing research programs along these lines will also help contextualize anthrax dynamics in the broader setting of global change biology. For example, a recent heat wave passing through Siberia released at least three separate outbreaks of *B. anthracis* preserved in now-melting permafrost. Our growing understanding of anthrax’s surprisingly long-term persistence indicates situations like this may become increasingly common in a changing world, and points towards the importance of linking reindeer herding practices to existing distributional patterns of anthrax spores and local wildlife communities. Similarly, the frameworks that have been used to study *B. anthracis* in Etosha could be invaluable to develop a research program assessing how widespread *B. cereus* biovar *anthracis* is in West and Central Africa, the basic routes of its transmission (especially given evidence flies play a strong role), and the magnitude of a threat it poses to conservation, agriculture, and human health. With existing work suggesting chimpanzee mortality might be so high that anthrax-like disease could effectively lead to their local extinction in the next century^26,114^, this work is just as pressing as work on classical *Bacillus anthracis.*

## III. Conclusions

In the face of global change, hidden rules that have produced extant landscapes of disease (on which theories are based) are liable to change, producing patterns that current interdisciplinary syntheses will sometimes fail to anticipate. In the face of these accelerating threats, we worry that the pace at which knowledge is collected and synthesized for pathogens like anthrax is not sufficient to keep pace, even with emerging interdisciplinary frameworks like One Health. The perspective on anthrax dynamics we present here has been loosely modeled off previous work on cholera, which has been widely noted as a model for One Health, interdisciplinarity, and the value of ecologists’ involvement in global health. We propose the long history of anthrax research can be a similar template for the value of multidisciplinary, investigative research. Anthrax is an especially useful model for the questions and complexities surrounding the transmission of environmentally-transmitted pathogens like brucellosis or plague, but we also note that the history of changing scientific knowledge about anthrax has broader lessons for pandemic prediction and prevention.

### (1) Lessons from Anthrax for Studying Environmental Transmission

On a broader scale, the seminal novelty of our multi-decade work on anthrax is a deeper understanding of how environmental maintenance and transmission affect the biology of a complex, multi-host pathogen. This has substantial relevance to research on other pathogens, including some of the most serious threats to human health; while some diseases like anthrax are primarily characterized by environmental modes of transmission, a far greater diversity of diseases are occasionally maintained in fomites and reservoirs. Many other bacteria appear to be capable of dormancy for extended periods of time, with no reproductive activity, within soil or aquatic environments. For example, *Clostridium botulinum* (the bacterium responsible for botulism) forms spores to persist in the environment, and many similar questions surround its biology, including whether spores germinate in unaffected hosts, including invertebrates and plants.^170^ Even familiar zoonotic pathogens with non-environmental routes of exposure can show surprising and unexpected patterns of environmental persistence. For example, recent work has confirmed that *Mycoplasma bovis*, a bacterial pathogen associated with mastitis in cattle and bison, can persist for long periods and possibly replicate, most likely through the formation of biofilms associated with gram negative bacteria, in sandy soil used as cattle bedding under certain moisture conditions.^171^ Similarly, plague (*Yersinia pestis)*, conventionally studied as a vector-borne disease, has been recently shown to persist in the soil for weeks—though the importance of this to the ecology and epidemiology of plague is still controversial.^172^

The questions we have highlighted here apply to any of these systems, and highlight key uncertainties in the role environmental transmission plays in these pathogens’ life cycles. In cases like these, the role that soil microbiota play in the dynamics and duration of persistence is predominantly unexplored and could represent a key future research direction. Similarly, specific environmental conditions, such as soil alkalinity, moisture, or specific mineral content, are required to allow environmental maintenance (possibly not through direct toxicity to the spores but through shaping the microbial community they interact with)—but for some systems, such as plague, those factors are still understudied or entirely untested. Ecological niche modeling tools that have been used to map anthrax persistence could easily be applied to other soil-borne bacteria, to elucidate the role of different drivers in persistence landscapes, and extrapolate transmission risk from microbiological knowledge.

From an eco-epidemiological angle, the role of environmental transmission in pathogen dynamics remains understudied and rarely modeled, especially in the case of diseases for which environmental transmission is not the primary mode of transmission.^173^ Key epidemiological concepts like *R_0_* rarely have the ability to accommodate environmental transmission, especially for generalist bacterial pathogens like anthrax.^174,175^ Environmental maintenance has a substantial effect on epidemiological dynamics even when a small part of a pathogen life cycle; for instance, Lowell *et al.* demonstrated that widespread plague epizootics are driven by local persistence in the soil for up to weeks at a time, a finding that can inform anticipatory surveillance of local factors (e.g. climate) known to increase plague outbreak risk.^172^ As many pathogens like plague utilize several transmission strategies, and may not rely primarily on environmental transmission (see Figure 1), modeling efforts for many pathogens may miss important dynamics if they exclude the occasional environmental route.^137,176–178^

Unusual evolutionary consequences of environmental transmission are also especially important. Evolutionary theory suggests that environmental persistence can release pathogens from virulence constraints^179^, something likely facilitated by broad generalism across diverse taxon groups. Two major fungal panzootics—chytrid (*Batrachochytrium dendrobatidis*) in amphibians ^180,181^ and white nose syndrome (*Pseudogymnoascus destructans*) in bats ^182,183^—are environmentally transmitted and maintained, and like anthrax and *B. cereus* biovar *anthracis*, are highly virulent generalist pathogens. Though neither fungus is zoonotic, the rapid emergence of these diseases poses an ongoing and worrisome problem for conservation efforts. On a different evolutionary timescale, just as pathogenic *B. cereus* outbreaks can be driven by long-term genetic cross-talk with *B. anthracis*, horizontal gene transfer from environmental reservoir strains can cause more abrupt changes in human pathogenic strains of other species. For example, in cholera, virulence genes transferred between benign and pathogenic strains in aquatic reservoirs are hypothesized to be the cause of some epidemics.^184^ Classical O1 and O139 strains and the El Tor (ET) strain of the 7^th^ pandemic appear to share common genetic elements like the *ctxA* and *tcpA* ET genes; genetic similarities between new toxigenic strains and these familiar strains are used to help understand the threat posed by new or unfamiliar strains^185^, and genetic surveillance in reservoirs has been proposed as a potential tool for outbreak anticipation.^185–187^

While bacterial and macroparasitic diseases more commonly evolve environmental modes of transmission, environmental persistence also plays an important role in the spatial epidemiology of some viruses, such as Hendra virus, for which viral shedding locations become spillover sites between bats and horses.^188^ Influenza can persist in waterways, and just like regulatory cross-talk has been important for anthrax and cholera outbreaks, strains of influenza that circulate in aquatic reservoirs and wild birds are likely to contribute new pathogenic strains to domestic poultry and potentially humans.^189^ Similarly, strains of polio can be transmitted in the environment from the live oral polio vaccine (OPV), leading to recurrent environmentally transmitted outbreaks that reinitiate polio transmission.^190^ Even many prionic diseases have an environmental transmission mode, such as scrapie^191^ or chronic wasting disease^174^, which persist in the soil for unknown durations, historically making pasture unusable for decades. As prionic diseases continue to spread through wild ungulates, and as grazing land infringes on natural areas, ecologists will be tasked with identifying and tracking agriculturally unsuitable, prion-contaminated land. If prions bond differently to different soil or vegetation types (as they appear to do ^192,193^), or host heterogeneity and movement determine the distribution of environmental reservoirs, the same One Health approaches that have succeeded in tracking anthrax emergence^169^ will need to be applied to the challenging problem of prion surveillance.

### (2) Lessons from Anthrax for Integrative Thinking

On a broader scale, our case study highlights the need for targeted interdisciplinarity in disease ecology. Interdisciplinary work, especially at long-term ecological research sites, has the potential to revise key ideas about pathogen biology and illuminate the hidden dynamics of pathogens in the environment. Some pathogens, like plague and cholera, are now well enough understood that forecasting can be done successfully across scales, ranging from local early warning systems to global projections under climate change.^15,194–197^ Anthrax poses a comparatively more serious challenge especially at broader scales, as locally developed scientific understanding becomes less transferrable; thus, expanding research across ecosystems with different dynamics and local drivers to extract generalities of environmental transmission dynamics represents a key next direction for synthesis work. But the majority of threats to public health are nowhere near that stage of synthesis. Ebola virus’s reservoirs are still uncertain, and the drivers of Ebola outbreaks have been recently studied but remain controversial at best.^198–200^ The enzootic cycle of Zika is even more poorly studied; the role primate reservoirs play in the enzootic process has been the subject of some speculation, but the ecology of the disease in its native range (Africa and potentially south Asia) remains essentially undocumented.^201^ The scope of complexity inherent to these pathogens’ life cycles cannot be fully understood until the enzootic process is better studied; and the ongoing value of interdisciplinarity as a tool for organizing that research is clear. We caution against a focus on research that pushes the cutting edges of disparate fields in isolation, which risks overlooking important insights gained from tying disparate fields together, and could leave the task of synthesis to policy makers with little or no scientific training. In the face of global change, interdisciplinary research is the only option for more rapid advances that keep pace with the accelerating threats that public health must face.

## IV. Acknowledgements

Eric Dougherty, Thomas Haverkamp, Jo Skeie Hermansen, and Hannah E. Schønhaug are gratefully acknowledged for their critical input on the concepts and writing of the manuscript. Dana Seidel is gratefully acknowledged for assistance with graphic design. All authors contributed to the design and main text of the paper. This work was supported by the National Socio-Environmental Synthesis Center (SESYNC) under funding received from the National Science Foundation DBI-1639145, the UC Berkeley Peder Sather Center for Advanced Study, The Research Council of Norway for core funding to CEES, and by NIH grant GM083863.

## References

1. Manlove, K. R. et al. ‘One Health’ or Three? Publication Silos Among the One Health Disciplines. PLoS Biol 14, e1002448 (2016).

2. Coker, R. et al. Towards a conceptual framework to support one-health research for policy on emerging zoonoses. Lancet Infect. Dis. 11, 326–331 (2011).

3. Parkes, M. W. et al. All hands on deck: transdisciplinary approaches to emerging infectious disease. EcoHealth 2, 258–272 (2005).

4. Borer, E. T. et al. Bridging taxonomic and disciplinary divides in infectious disease. EcoHealth 8, 261–267 (2011).

5. Jones, K. E. et al. Global trends in emerging infectious diseases. Nature 451, 990–993 (2008).

6. Johnson, P. T., De Roode, J. C. & Fenton, A. Why infectious disease research needs community ecology. Science 349, 1259504 (2015).

7. Ezenwa, V. O. et al. Interdisciplinarity and infectious diseases: An ebola case study. PLoS Pathog 11, e1004992 (2015).

8. Wilcox, B. A. & Colwell, R. R. Emerging and reemerging infectious diseases: biocomplexity as an interdisciplinary paradigm. EcoHealth 2, 244 (2005).

9. Wilcox, B. A. & Gubler, D. J. Disease ecology and the global emergence of zoonotic pathogens. Environ. Health Prev. Med. 10, 263–272 (2005).

10. Daszak, P. et al. Collaborative research approaches to the role of wildlife in zoonotic disease emergence. in Wildlife and emerging zoonotic diseases: the biology, circumstances and consequences of cross-species transmission 463–475 (Springer, 2007).

11. Levin, B. R., Lipsitch, M. & Bonhoeffer, S. Population biology, evolution, and infectious disease: convergence and synthesis. Science 283, 806–809 (1999).

12. Moore, C. G. Interdisciplinary research in the ecology of vector-borne diseases: Opportunities and needs. J. Vector Ecol. 33, 218–224 (2008).

13. Plowright, R. K., Sokolow, S. H., Gorman, M. E., Daszak, P. & Foley, J. E. Causal inference in disease ecology: investigating ecological drivers of disease emergence. Front. Ecol. Environ. 6, 420–429 (2008).

14. Huq, A. et al. Ecological relationships between Vibrio cholerae and planktonic crustacean copepods. Appl. Environ. Microbiol. 45, 275–283 (1983).

15. Colwell, R. R. Global climate and infectious disease: the cholera paradigm. Science 274, 2025 (1996).

16. Colwell, R. R. et al. Reduction of cholera in Bangladeshi villages by simple filtration. Proc. Natl. Acad. Sci. 100, 1051–1055 (2003).

17. Hugh-Jones, M. & De Vos, V. Anthrax and wildlife. Rev. Sci. Tech.-Off. Int. Epizoot. 21, 359–384 (2002).

18. Hugh-Jones, M. & Blackburn, J. The ecology of Bacillus anthracis. Mol. Aspects Med. 30, 356–367 (2009).

19. Turner, W. C. et al. Fatal attraction: vegetation responses to nutrient inputs attract herbivores to infectious anthrax carcass sites. Proc. R. Soc. Lond. B Biol. Sci. 281, 20141785 (2014).

20. Blackburn, J. K., Van Ert, M., Mullins, J. C., Hadfield, T. L. & Hugh-Jones, M. E. The necrophagous fly anthrax transmission pathway: empirical and genetic evidence from wildlife epizootics. Vector-Borne Zoonotic Dis. 14, 576–583 (2014).

21. Barandongo, Z. R., Mfune, J. K. & Turner, W. C. Dust-bathing behaviors of African herbivores and the potential risk of inhalation anthrax. J. Wildl. Dis. 54, 34–44 (2018).

22. Turner, W. C. et al. Lethal exposure: An integrated approach to pathogen transmission via environmental reservoirs. Sci. Rep. 6, (2016).

23. Shadomy, S. V. & Smith, T. L. Anthrax. J. Am. Vet. Med. Assoc. 233, 63–72 (2008).

24. Beatty, M. E., Ashford, D. A., Griffin, P. M., Tauxe, R. V. & Sobel, J. Gastrointestinal anthrax: review of the literature. Arch. Intern. Med. 163, 2527–2531 (2003).

25. Bales, M. E. et al. Epidemiologic responses to anthrax outbreaks: a review of field investigations, 1950-2001. Emerg. Infect. Dis. 8, 1163 (2002).

26. Hoffmann, C. et al. Persistent anthrax as a major driver of wildlife mortality in a tropical rainforest. Nature 548, 82–86 (2017).

27. Lindeque, P. & Turnbull, P. Ecology and epidemiology of anthrax in the Etosha National Park, Namibia. Onderstepoort J Vet Res 61, 71–83 (1994).

28. Antonation, K. S. et al. Bacillus cereus biovar anthracis causing anthrax in sub-Saharan Africa—chromosomal monophyly and broad geographic distribution. PLoS Negl. Trop. Dis. 10, e0004923 (2016).

29. Debré, P. Louis Pasteur. (Johns Hopkins University Press, 2000).

30. Manchee, R., Broster, M., Melling, J., Henstridge, R. & Stagg, A. Bacillus anthracis on Gruinard island. Nature 294, 254–255 (1981).

31. Williams, G., Linley, E., Nicholas, R. & Baillie, L. The role of the exosporium in the environmental distribution of anthrax. J. Appl. Microbiol. 114, 396–403 (2013).

32. Turnbull, P. et al. Bacillus anthracis but not always anthrax. J. Appl. Microbiol. 72, 21–28 (1992).

33. Braun, P. et al. Microevolution of anthrax from a young ancestor (MAYA) suggests a soil-borne life cycle of Bacillus anthracis. PloS One 10, e0135346 (2015).

34. Van Ness, G. B. Ecology of anthrax. Science 172, 1303–1307 (1971).

35. Minett, F. & Dhanda, M. Multiplication of B. anthracis and Cl. chauvei in soil and water. Ind J Vet Sci Anim Husb 11, 308–328 (1941).

36. Minett, F. Sporulation and viability of B. anthracis in relation to environmental temperature and humidity. J. Comp. Pathol. Ther. 60, 161–176 (1950).

37. Vasil’eva, V. Soil bacteria as antagonists of anthrax bacilli. Vet. Bull. 9, 149–153 (1958).

38. Zarubkinskii, V. Self purification of soil and water from anthrax bacilli. Vet. Bull. 9, 51–58 (1958).

39. Hendriksen, N. B. & Carstensen, J. Long-term survival of Bacillus thuringiensis subsp. kurstaki in a field trial. Can. J. Microbiol. 59, 34–38 (2013).

40. West, A., Burges, H., Dixon, T. & Wyborn, C. Survival of Bacillus thuringiensis and Bacillus cereus spore inocula in soil: effects of pH, moisture, nutrient availability and indigenous microorganisms. Soil Biol. Biochem. 17, 657–665 (1985).

41. Yara, K., Kunimi, Y. & Iwahana, H. Comparative studies of growth characteristic and competitive ability in Bacillus thuringiensis and Bacillus cereus in soil. Appl. Entomol. Zool. 32, 625–634 (1997).

42. Saile, E. & Koehler, T. M. Bacillus anthracis multiplication, persistence, and genetic exchange in the rhizosphere of grass plants. Appl. Environ. Microbiol. 72, 3168–3174 (2006).

43. Halverson, L. J., Clayton, M. K. & Handelsman, J. Population biology of Bacillus cereus UW85 in the rhizosphere of field-grown soybeans. Soil Biol. Biochem. 25, 485–493 (1993).

44. Raymond, B., Johnston, P. R., Nielsen-LeRoux, C., Lereclus, D. & Crickmore, N. Bacillus thuringiensis: an impotent pathogen? Trends Microbiol. 18, 189–194 (2010).

45. Monnerat, R. G. et al. Translocation and insecticidal activity of Bacillus thuringiensis living inside of plants. Microb. Biotechnol. 2, 512–520 (2009).

46. Vidal-Quist, J. C., Rogers, H. J., Mahenthiralingam, E. & Berry, C. Bacillus thuringiensis colonises plant roots in a phylogeny-dependent manner. FEMS Microbiol. Ecol. 86, 474–489 (2013).

47. Beauregard, P. B., Chai, Y., Vlamakis, H., Losick, R. & Kolter, R. Bacillus subtilis biofilm induction by plant polysaccharides. Proc. Natl. Acad. Sci. 110, E1621–E1630 (2013).

48. Chen, Y. et al. Biocontrol of tomato wilt disease by Bacillus subtilis isolates from natural environments depends on conserved genes mediating biofilm formation. Environ. Microbiol. 15, 848–864 (2013).

49. Lee, K. et al. Phenotypic and functional characterization of Bacillus anthracis biofilms. Microbiology 153, 1693–1701 (2007).

50. Schwartz, M. The life and works of Louis Pasteur. J. Appl. Microbiol. 91, 597–601 (2001).

51. Schuch, R. & Fischetti, V. A. The secret life of the anthrax agent Bacillus anthracis: bacteriophage-mediated ecological adaptations. PloS One 4, e6532 (2009).

52. Dey, R., Hoffman, P. S. & Glomski, I. J. Germination and amplification of anthrax spores by soil-dwelling amoebas. Appl. Environ. Microbiol. 78, 8075–8081 (2012).

53. Barbaree, J. M., Fields, B. S., Feeley, J. C., Gorman, G. W. & Martin, W. T. Isolation of protozoa from water associated with a legionellosis outbreak and demonstration of intracellular multiplication of Legionella pneumophila. Appl. Environ. Microbiol. 51, 422–424 (1986).

54. El-Etr, S. H. et al. Francisella tularensis type A strains cause the rapid encystment of Acanthamoeba castellanii and survive in amoebal cysts for three weeks postinfection. Appl. Environ. Microbiol. 75, 7488–7500 (2009).

55. Kolstø, A.-B., Tourasse, N. J. & Økstad, O. A. What sets Bacillus anthracis apart from other Bacillus species? Annu. Rev. Microbiol. 63, 451–476 (2009).

56. Helgason, E., Tourasse, N. J., Meisal, R., Caugant, D. A. & Kolstø, A.-B. Multilocus sequence typing scheme for bacteria of the Bacillus cereus group. Appl. Environ. Microbiol. 70, 191–201 (2004).

57. Zwick, M. E. et al. Genomic characterization of the Bacillus cereus sensu lato species: backdrop to the evolution of Bacillus anthracis. Genome Res. 22, 1512–1524 (2012).

58. Priest, F. G., Barker, M., Baillie, L. W., Holmes, E. C. & Maiden, M. C. Population structure and evolution of the Bacillus cereus group. J. Bacteriol. 186, 7959–7970 (2004).

59. Bazinet, A. L. Pan-genome and phylogeny of Bacillus cereus sensu lato. BMC Evol. Biol. 17, 176 (2017).

60. Zheng, J. et al. Comparative genomics of Bacillus thuringiensis reveals a path to specialized exploitation of multiple invertebrate hosts. MBio 8, e00822–17 (2017).

61. Pilo, P. & Frey, J. Bacillus anthracis: Molecular taxonomy, population genetics, phylogeny and patho-evolution. Infect. Genet. Evol. 11, 1218–1224 (2011).

62. Keim, P. et al. Anthrax molecular epidemiology and forensics: using the appropriate marker for different evolutionary scales. Infect. Genet. Evol. 4, 205–213 (2004).

63. Van Ert, M. N. et al. Global genetic population structure of Bacillus anthracis. PloS One 2, e461 (2007).

64. Beyer, W. et al. Distribution and molecular evolution of Bacillus anthracis genotypes in Namibia. PLoS Negl. Trop. Dis. 6, e1534 (2012).

65. Beyer, W. & Turnbull, P. Co-infection of an animal with more than one genotype can occur in anthrax. Lett. Appl. Microbiol. 57, 380–384 (2013).

66. Kenefic, L. J. et al. Pre-columbian origins for north american anthrax. PLoS One 4, e4813 (2009).

67. Hoffmaster, A. R. et al. Identification of anthrax toxin genes in a Bacillus cereus associated with an illness resembling inhalation anthrax. Proc. Natl. Acad. Sci. U. S. A. 101, 8449–8454 (2004).

68. Leendertz, F. H. et al. Anthrax kills wild chimpanzees in a tropical rainforest. Nature 430, 451–452 (2004).

69. Klee, S. R. et al. Characterization of Bacillus anthracis-like bacteria isolated from wild great apes from Cote d’Ivoire and Cameroon. J. Bacteriol. 188, 5333–5344 (2006).

70. Klee, S. R. et al. The genome of a Bacillus isolate causing anthrax in chimpanzees combines chromosomal properties of B. cereus with B. anthracis virulence plasmids. PloS One 5, e10986 (2010).

71. Brézillon, C. et al. Capsules, toxins and AtxA as virulence factors of emerging Bacillus cereus biovar anthracis. PLoS Negl. Trop. Dis. 9, e0003455 (2015).

72. Oh, S.-Y., Budzik, J. M., Garufi, G. & Schneewind, O. Two capsular polysaccharides enable Bacillus cereus G9241 to cause anthrax-like disease. Mol. Microbiol. 80, 455–470 (2011).

73. Leppla, S. H. Anthrax Toxin Edema Factor: A Bacterial Adenylate Cyclase That Increases Cyclic AMP Concentrations in Eukaryotic Cells. Proc. Natl. Acad. Sci. 79, 3162–3166 (1982).

74. Collier, R. J. & Young, J. A. T. Anthrax Toxin. Annu Rev Cell Dev Biol 19, 45–70 (2003).

75. Brien, J. O., Friedlander, A., Dreier, T., Ezzell, J. & Leppla, S. Effects of Anthrax Toxin Components on Human Neutrophils. Infect. Immun. 47, 306–310 (1985).

76. Comer, J. E., Chopra, A. K., Peterson, J. W. & Konig, R. Direct Inhibition of T- Lymphocyte Activation by Anthrax Toxins In Vivo. Infect. Immun. 73, 8275–8281 (2005).

77. Duesbery, N. S. Proteolytic Inactivation of MAP-Kinase-Kinase by Anthrax Lethal Factor. Science 280, 734–737 (1998).

78. Vitale, G. et al. Anthrax Lethal Factor Cleaves the N-Terminus of MAPKKs and Induces Tyrosine/Threonine Phosphorylation of MAPKs in Cultured Macrophages. Biochem. Biophys. Res. Commun. 248, 706–711 (1998).

79. Erwin, J. L. et al. Macrophage-Derived Cell Lines Do Not Express Proinflammatory Cytokines after Exposure to Bacillus anthracis Lethal Toxin. Infect. Immun. 69, 1175–1177 (2001).

80. Pellizzari, R., Guidi-Rontani, C., Vitale, G., Mock, M. & Montecucco, C. Anthrax lethal factor cleaves MKK3 in macrophages and inhibits the LPS/IFNgamma-induced release of NO and TNFalpha. FEBS Lett. 462, 199–204 (1999).

81. Ribot, W. J. et al. Anthrax lethal toxin impairs innate immune functions of alveolar macrophages and facilitates Bacillus anthracis survival. Infect. Immun. 74, 5029–34 (2006).

82. Agrawal, A. et al. Impairment of dendritic cells and adaptive immunity by anthrax lethal toxin. Nature 424, 329–34 (2003).

83. Mogridge, J., Cunningham, K. & Collier, R. J. Stoichiometry of anthrax toxin complexes. Biochemistry (Mosc.) 41, 1079–1082 (2002).

84. Fink, S. L., Bergsbaken, T. & Cookson, B. T. Anthrax lethal toxin and Salmonella elicit the common cell death pathway of caspase-1-dependent pyroptosis via distinct mechanisms. Proc. Natl. Acad. Sci. 105, 4312–4317 (2008).

85. Aloni-Grinstein, R. et al. Oral Spore Vaccine Based on Live Attenuated Nontoxinogenic Bacillus anthracis Expressing Recombinant Mutant Protective Antigen. Infect. Immun. 73, 4043–4053 (2005).

86. Cohen, S. N. et al. Attenuated Nontoxinogenic and Nonencapsulated Recombinant Bacillus anthracis Spore Vaccines Protect against Anthrax. Infect. Immun. 68, 4549–4558 (2000).

87. Little, S. F., Ivins, B. E., Fellows, P. F. & Fried. Passive protection by polyclonal antibodies against Bacillus anthracis Infection in guinea pigs. Infect. Immun. 65, 5171–5175 (1997).

88. Little, S. F. et al. Development of an in vitro-based potency assay for anthrax vaccine. Vaccine 22, 2843–2852 (2004).

89. Marcus, H. et al. Contribution of immunological memory to protective immunity conferred by Bacillus anthracis protective antigen-based vaccine. Infect. Immun. 72, 3471–3477 (2004).

90. Pitt, M. L. M. et al. In vitro correlate of immunity in a rabbit model of inhalational anthrax. Appl. Microbiol. 19, 4768–4773 (2001).

91. Reuveny, S. et al. Search for correlates of protective immunity conferred by anthrax vaccine. Infect. Immun. 69, 2888–2893 (2001).

92. Turnbull, P.C. B. Current status of immunization against anthrax: Old vaccines may be here to stay for a while. Curr. Opin. Infect. Dis. 13, 113–120 (2000).

93. Allen, J. S., Skowera, A., Rubin, G. J., Wessely, S. & Peakman, M. Long-lasting T cell responses to biological warfare vaccines in human vaccinees. Clin. Infect. Dis. 43, 1–7 (2006).

94. Cote, C. K., Rea, K. M., Norris, S. L., van Rooijen, N. & Welkos, S. L. The use of a model of in vivo macrophage depletion to study the role of macrophages during infection with Bacillus anthracis spores. Microb. Pathog. 37, 169–175 (2004).

95. Cote, C. K., Van Rooijen, N. & Welkos, S. L. Roles of macrophages and neutrophils in the early host response to Bacillus anthracis spores in a mouse model of infection. Infect. Immun. 74, 469–480 (2006).

96. Barton, G. M. & Medzhitov, R. Toll-like receptor signaling pathways. Science 300, 1524–1525 (2003).

97. Yadav, J. S. et al. Multigenic control and sex bias in host susceptibility to spore-induced pulmonary anthrax in mice. Infect. Immun. 79, 3204–3215 (2011).

98. Hughes, M. A. et al. MyD88-dependent signaling contributes to protection following Bacillus anthracis spore challenge of mice: implications for Toll-like receptor signaling. Infect. Immun. 73, 7535–7540 (2005).

99. Boyden, E. D. & Dietrich, W. F. Nalp1b controls mouse macrophage susceptibility to anthrax lethal toxin. Nat. Genet. 38, 240–244 (2006).

100. Liao, K.-C. & Mogridge, J. Expression of Nlrp1b inflammasome components in human fibroblasts confers susceptibility to anthrax lethal toxin. Infect. Immun. 77, 4455–4462 (2009).

101. Terra, J. K. et al. Cutting edge: resistance to Bacillus anthracis infection mediated by a lethal toxin sensitive allele of Nalp1b/Nlrp1b. J. Immunol. 184, 17–20 (2010).

102. Moayeri, M. et al. Inflammasome sensor Nlrp1b-dependent resistance to anthrax is mediated by caspase-1, IL-1 signaling and neutrophil recruitment. PLoS Pathog 6, e1001222 (2010).

103. Bradley, K. A., Mogridge, J., Mourez, M., Collier, R. J. & Young, J. A. Identification of the cellular receptor for anthrax toxin. Nature 414, 225–229 (2001).

104. Scobie, H. M., Rainey, G. J. A., Bradley, K. A. & Young, J. A. Human capillary morphogenesis protein 2 functions as an anthrax toxin receptor. Proc. Natl. Acad. Sci. 100, 5170–5174 (2003).

105. Liu, S. et al. Capillary morphogenesis protein-2 is the major receptor mediating lethality of anthrax toxin in vivo. Proc. Natl. Acad. Sci. 106, 12424–12429 (2009).

106. Martchenko, M., Candille, S. I., Tang, H. & Cohen, S. N. Human genetic variation altering anthrax toxin sensitivity. Proc. Natl. Acad. Sci. 109, 2972–2977 (2012).

107. Turnbull, P. C. B., Doganay, M., Lindeque, P. M., Aygen, B. & McLaughlin, J. Serology and anthrax in humans, livestock and Etosha National Park wildlife. Epidemiol. Infect. 108, 299–313 (1992).

108. Turnbull, P. C. B. et al. Naturally acquired antibodies to Bacillus anthracis protective antigen in vultures in southern Africa. Onderstepoort J. Vet. Res. 75, 95–102 (2008).

109. Lembo, T. et al. Serologic Surveillance of Anthrax in the Serengeti Ecosystem, Tanzania, 1996-2009. Emerg. Infect. Dis. 17, 387–394 (2011).

110. Hampson, K. et al. Predictability of anthrax infection in the Serengeti, Tanzania. J. Appl. Ecol. no-no (2011). doi:10.1111/j.1365-2664.2011.02030.x

111. Bellan, S. E. et al. Black-backed jackal exposure to rabies virus, canine distemper virus, and bacillus anthracis in etosha national park, namibia. J. Wildl. Dis. 48, 371–81 (2012).

112. Cizauskas, C. A., Bellan, S. E., Turner, W. C., Vance, R. E. & Getz, W. M. Frequent and seasonally variable sublethal anthrax infections are accompanied by short-lived immunity in an endemic system. J. Anim. Ecol. 83, 1078–1090 (2014).

113. Caraco, T. & Turner, W. C. Pathogen transmission at stage-structured infectious patches: Killers and vaccinators. J. Theor. Biol. 436, 51–63 (2018).

114. Zimmermann, F. et al. Low antibody prevalence against Bacillus cereus biovar anthracis in Tai National Park, Cote d’Ivoire, indicates high rate of lethal infections in wildlife. PLoS Negl. Trop. Dis. 11, e0005960 (2017).

115. Cizauskas, C. A. et al. Gastrointestinal helminths may affect host susceptibility to anthrax through seasonal immune trade-offs. BMC Ecol. 14, 27 (2014).

116. Cizauskas, C. A., Turner, W. C., Pitts, N. & Getz, W. M. Seasonal patterns of hormones, macroparasites, and microparasites in wild African ungulates: the interplay among stress, reproduction, and disease. PloS One 10, e0120800 (2015).

117. Getz, W. M. Biomass transformation webs provide a unified approach to consumer-resource modelling. Ecol. Lett. 14, 113–124 (2011).

118. Ganz, H. H. et al. Interactions between Bacillus anthracis and plants may promote anthrax transmission. PLoSNegl Trop Dis 8, e2903 (2014).

119. Getz, W. M., Salter, R., Seidel, D. P. & Hooft, P. Sympatric speciation in structureless environments. BMC Evol. Biol. 16, 50 (2016).

120. Turner, W. C. et al. Soil ingestion, nutrition and the seasonality of anthrax in herbivores of Etosha National Park. Ecosphere

121. Dougherty, E. R., Seidel, D. P., Carlson, C. J. & Getz, W. M. Using movement data to estimate contact rates in a simulated environmentally-transmitted disease system. bioRxiv 261198 (2018).

122. Dougherty, E. R., Carlson, C. J., Blackburn, J. K. & Getz, W. M. A cross-validation-based approach for delimiting reliable home range estimates. Mov. Ecol. 5, 19 (2017).

123. Zidon, R., Garti, S., Getz, W. M. & Saltz, D. Zebra migration strategies and anthrax in Etosha National Park, Namibia. Ecosphere

124. Mashintonio, A. F., Pimm, S. L., Harris, G. M., Van Aarde, R. J. & Russell, G. J. Data-driven discovery of the spatial scales of habitat choice by elephants. PeerJ 2, e504 (2014).

125. Lyons, A. J., Turner, W. C. & Getz, W. M. Home range plus: a space-time characterization of movement over real landscapes. Mov. Ecol. 1, 2 (2013).

126. Polansky, L., Kilian, W. & Wittemyer, G. Elucidating the significance of spatial memory on movement decisions by African savannah elephants using state-space models. in Proc. R. Soc. B 282, 20143042 (The Royal Society, 2015).

127. Morris, L. R., Proffitt, K. M., Asher, V. & Blackburn, J. K. Elk resource selection and implications for anthrax management in Montana. J. Wildl. Manag. 80, 235–244 (2016).

128. Dougherty, E. R., Seidel, D. P., Carlson, C. J., Speigel, O. & Getz, W. M. Going through the motions: incorporating movement analyses into disease research. Ecol. Lett. in press (2018). doi:10.1111/ele.12917

129. Bullock, D. S. Vultures as disseminators of anthrax. The Auk 283–284 (1956).

130. Houston, D. C. & Cooper, J. The digestive tract of the whiteback griffon vulture and its role in disease transmission among wild ungulates. J. Wildl. Dis. 11, 306–313 (1975).

131. Urbain, A. & Novel, J. Spread of tuberculosis and anthrax by carnivorous birds. Bull Acad Vet Fr 19, 237–239 (1946).

132. Ganz, H. H., Karaoz, U., Getz, W. M., Versfeld, W. & Brodie, E. L. Diversity and structure of soil bacterial communities associated with vultures in an African savanna. Ecosphere 3, 1–18 (2012).

133. Davies, D. The influence of temperature and humidity on spore formation and germination in Bacillus anthracis. Epidemiol. Infect. 58, 177–186 (1960).

134. Bellan, S. E., Turnbull, P. C., Beyer, W. & Getz, W. M. Effects of experimental exclusion of scavengers from carcasses of anthrax-infected herbivores on Bacillus anthracis sporulation, survival, and distribution. Appl. Environ. Microbiol. 79, 3756–3761 (2013).

135. Getz, W. M. et al. Making ecological models adequate. Ecol. Lett. 21, 153–166 (2018).

136. Brouwer, A. F., Weir, M. H., Eisenberg, M. C., Meza, R. & Eisenberg, J. N. Dose-response relationships for environmentally mediated infectious disease transmission models. PLoS Comput. Biol. 13, e1005481 (2017).

137. McCallum, H. et al. Breaking beta: deconstructing the parasite transmission function. Phil Trans R Soc B 372, 20160084 (2017).

138. Turner, A., Galvin, J., Rubira, R. & Miller, G. Anthrax explodes in an Australian summer. J. Appl. Microbiol. 87, 196–199 (1999).

139. Parkinson, R., Rajic, A. & Jenson, C. Investigation of an anthrax outbreak in Alberta in 1999 using a geographic information system. Can. Vet. J. 44, 315 (2003).

140. Blackburn, J. K. & Goodin, D. G. Differentiation of springtime vegetation indices associated with summer anthrax epizootics in west Texas, USA, deer. J. Wildl. Dis. 49, 699–703 (2013).

141. Barro, A. S. et al. Redefining the Australian anthrax belt: Modeling the ecological niche and predicting the geographic distribution of Bacillus anthracis. PLoS Negl Trop Dis 10, e0004689 (2016).

142. Blackburn, J. K. et al. Bacillus anthracis diversity and geographic potential across Nigeria, Cameroon and Chad: further support of a novel West African lineage. PLoS Negl Trop Dis 9, e0003931 (2015).

143. Chen, W.-J. et al. Mapping the Distribution of Anthrax in Mainland China, 2005-2013. PLoS Negl Trop Dis 10, e0004637 (2016).

144. Kracalik, I. T. et al. Modeling the environmental suitability of anthrax in Ghana and estimating populations at risk: Implications for vaccination and control. PLoS Negl‥ Trop. Dis. 11, e0005885 (2017).

145. Mullins, J. C. et al. Ecological niche modeling of Bacillus anthracis on three continents: evidence for genetic-ecological divergence? PloS One 8, e72451 (2013).

146. Blackburn, J. K. et al. Modeling the Ecological Niche of Bacillus anthracis to Map Anthrax Risk in Kyrgyzstan. Am. J. Trop. Med. Hyg. 96, 550–556 (2017).

147. Blackburn, J. K. Integrating geographic information systems and ecological niche modeling into disease ecology: a case study of Bacillus anthracis in the United States and Mexico. in Emerging and Endemic Pathogens 59–88 (Springer, 2010).

148. Chikerema, S., Murwira, A., Matope, G. & Pfukenyi, D. Spatial modelling of Bacillus anthracis ecological niche in Zimbabwe. Prev. Vet. Med. 111, 25–30 (2013).

149. Joyner, T. A. et al. Modeling the potential distribution of Bacillus anthracis under multiple climate change scenarios for Kazakhstan. PloS One 5, e9596 (2010).

150. Walsh, M. G., de Smalen, A. W. & Mor, S. Climatic influence on the anthrax niche in warming northern latitudes. bioRxiv 219857 (2017).

151. Iacono, G. L. et al. A unified framework for the infection dynamics of zoonotic spillover and spread. PLoS Negl. Trop. Dis. 10, e0004957 (2016).

152. Redding, D. W., Moses, L. M., Cunningham, A. A., Wood, J. & Jones, K. E. Environmental-mechanistic modelling of the impact of global change on human zoonotic disease emergence: a case study of Lassa fever. Methods Ecol. Evol. 7, 646–655 (2016).

153. Redding, D. et al. Impact of global change on future Ebola emergence and epidemic potential in Africa. bioRxiv 206169 (2017).

154. Brett, T. S., Drake, J. M. & Rohani, P. Anticipating the emergence of infectious diseases. J. R. Soc. Interface 14, 20170115 (2017).

155. O’Regan, S. M. & Drake, J. M. Theory of early warning signals of disease emergenceand leading indicators of elimination. Theor. Ecol. 6, 333–357 (2013).

156. Thomson, M. et al. Malaria early warnings based on seasonal climate forecasts from multi-model ensembles. Nature 439, 576 (2006).

157. Semenza, J. C. et al. Environmental Suitability of Vibrio Infections in a Warming Climate: An Early Warning System. Environ. Health Perspect. 125, (2017).

158. Bellan, S. E., Gimenez, O., Choquet, R. & Getz, W. M. A hierarchical distance sampling approach to estimating mortality rates from opportunistic carcass surveillance data. Methods Ecol. Evol. 4, 361–369 (2013).

159. Opare, C., Nsiire, A., Awumbilla, B. & Akanmori, B. Human behavioural factors implicated in outbreaks of human anthrax in the Tamale municipality of northern Ghana. Acta Trop. 76, 49–52 (2000).

160. Gainer, R. & Oksanen, A. Anthrax and the Taiga. Can. Vet. J. 53, 1123 (2012).

161. Unnamed reporter. Two more outbreaks of anthrax hit northern Siberia due to thawing permafrost. The Siberian Times (2016).

162. Unnamed reporter. UPDATED First anthrax outbreak since 1941: 9 hospitalised, with two feared to have disease. The Siberian Times (2016).

163. Gertcyk, O. Huge cull of 250,000 reindeer by Christmas in Yamalo-Nenets after anthrax outbreak. The Siberian Times (2016).

164. Doucleff, M. Killing Reindeer To Stop Anthrax Could Snuff Out A Nomadic Culture. NPR (2016).

165. Schmid, G. & Kaufmann, A. Anthrax in Europe: its epidemiology, clinical characteristics, and role in bioterrorism. Clin. Microbiol. Infect. 8, 479–488 (2002).

166. Unknown. Anthrax outbreak kills nine animals in Sweden. The Local (2016).

167. Blackburn, J. K., McNyset, K. M., Curtis, A. & Hugh-Jones, M. E. Modeling the geographic distribution of Bacillus anthracis, the causative agent of anthrax disease, for the contiguous United States using predictive ecologic niche modeling. Am. J. Trop. Med. Hyg. 77, 1103–1110 (2007).

168. Ebedes, H. Anthrax epizootics in Etosha National Park. Madoqua 10, 99–118 (1977).

169. Blackburn, J. K., Kracalik, I. T. & Fair, J. M. Applying Science: Opportunities to Inform Disease Management Policy with Cooperative Research within a One Health Framework. Front. Public Health 3, (2015).

170. Espelund, M. & Klaveness, D. Botulism outbreaks in natural environments-an update. Front. Microbiol. 5, 287 (2014).

171. Justice-Allen, A. et al. Survival and replication of Mycoplasma species in recycled bedding sand and association with mastitis on dairy farms in Utah. J. Dairy Sci. 93, 192–202 (2010).

172. Lowell, J. L. et al. Single-nucleotide polymorphisms reveal spatial diversity among clones of Yersinia pestis during plague outbreaks in Colorado and the western United States. Vector-Borne Zoonotic Dis. 15, 291–302 (2015).

173. Saad-Roy, C., van den Driessche, P. & Yakubu, A.-A. A Mathematical Model of Anthrax Transmission in Animal Populations. Bull. Math. Biol. 79, 303–324 (2017).

174. Almberg, E. S., Cross, P. C., Johnson, C. J., Heisey, D. M. & Richards, B. J. Modeling Routes of Chronic Wasting Disease Transmission: Environmental Prion Persistence Promotes Deer Population Decline and Extinction. PLOS ONE 6, 1–11 (2011).

175. Ivanek, R. & Lahodny Jr, G. From the bench to modeling-R0 at the interface between empirical and theoretical approaches in epidemiology of environmentally transmitted infectious diseases. Prev. Vet. Med. 118, 196–206 (2015).

176. Mata, M. A. E., Greenwood, P. E. & Tyson, R. C. The roles of direct and environmental transmission in stochastic avian flu epidemic recurrence. ArXiv Prepr. ArXiv170304869 (2017).

177. Lange, M., Kramer-Schadt, S. & Thulke, H.-H. Relevance of Indirect Transmission for Wildlife Disease Surveillance. Front. Vet. Sci. 3, 110 (2016).

178. Tien, J. H. & Earn, D. J. Multiple transmission pathways and disease dynamics in a waterborne pathogen model. Bull. Math. Biol. 72, 1506–1533 (2010).

179. Roche, B., Drake, J. M. & Rohani, P. The curse of the Pharaoh revisited: evolutionary bistability in environmentally transmitted pathogens. Ecol. Lett. 14, 569–575 (2011).

180. Kilpatrick, A. M., Briggs, C. J. & Daszak, P. The ecology and impact of chytridiomycosis: an emerging disease of amphibians. Trends Ecol. Evol. 25, 109–118 (2010).

181. Mitchell, K. M., Churcher, T. S., Garner, T. W. & Fisher, M. C. Persistence of the emerging pathogen Batrachochytrium dendrobatidis outside the amphibian host greatly increases the probability of host extinction. Proc. R. Soc. Lond. B Biol. Sci. 275, 329–334 (2008).

182. Lorch, J. M. et al. Distribution and environmental persistence of the causative agent of white-nose syndrome, Geomyces destructans, in bat hibernacula of the eastern United States. Appl. Environ. Microbiol. 79, 1293–1301 (2013).

183. Reynolds, H. T., Ingersoll, T. & Barton, H. A. Modeling the environmental growth of Pseudogymnoascus destructans and its impact on the white-nose syndrome epidemic. J. Wildl. Dis. 51, 318–331 (2015).

184. Faruque, S. M. & Nair, G. B. Molecular ecology of toxigenic Vibrio cholerae. Microbiol. Immunol. 46, 59–66 (2002).

185. Bik, E. M., Gouw, R. D. & Mooi, F. R. DNA fingerprinting of Vibrio cholerae strains with a novel insertion sequence element: a tool to identify epidemic strains. J. Clin. Microbiol. 34, 1453–1461 (1996).

186. De, R., Ghosh, J. B., Sen Gupta, S., Takeda, Y. & Nair, G. B. The role of Vibrio cholerae genotyping in Africa. J. Infect. Dis. 208, S32–S38 (2013).

187. Rivera, I. N., Chun, J., Huq, A., Sack, R. B. & Colwell, R. R. Genotypes associated with virulence in environmental isolates of Vibrio cholerae. Appl. Environ. Microbiol. 67, 2421–2429 (2001).

188. Plowright, R. K. et al. Ecological dynamics of emerging bat virus spillover. in Proc. R. Soc. B 282, 20142124 (The Royal Society, 2015).

189. Ip, H. S. et al. Novel Eurasian Highly Pathogenic Avian Influenza A H5 Viruses in Wild Birds, Washington, USA, 2014. Emerg. Infect. Dis. J. 21, 886 (2015).

190. Fine, P. E. & Carneiro, I. A. Transmissibility and persistence of oral polio vaccine viruses: implications for the global poliomyelitis eradication initiative. Am. J. Epidemiol. 150, 1001–1021 (1999).

191. Seidel, B. et al. Scrapie agent (strain 263K) can transmit disease via the oral route after persistence in soil over years. PLoS One 2, e435 (2007).

192. Johnson, C. J., Pedersen, J. A., Chappell, R. J., McKenzie, D. & Aiken, J. M. Oral transmissibility of prion disease is enhanced by binding to soil particles. PLoS Pathog. 3, e93 (2007).

193. Pritzkow, S. et al. Grass plants bind, retain, uptake, and transport infectious prions. Cell Rep. 11, 1168–1175 (2015).

194. Bramanti, B., Stenseth, N. C., Walløe, L. & Lei, X. Plague: A Disease Which Changed the Path of Human Civilization. in Yersinia pestis: Retrospective and Perspective 1–26 (Springer, 2016).

195. Stenseth, N. C. et al. Plague dynamics are driven by climate variation. Proc. Natl. Acad. Sci. 103, 13110–13115 (2006).

196. Stenseth, N. C. et al. Plague: past, present, and future. PLoS Med. 5, e3 (2008).

197. Escobar, L. E. et al. A global map of suitability for coastal Vibrio cholerae under current and future climate conditions. Acta Trop. 149, 202–211 (2015).

198. Alexander, K. A. et al. What factors might have led to the emergence of Ebola in West Africa? PLoS Negl. Trop. Dis. 9, e0003652 (2015).

199. Wallace, R. G. et al. Did Ebola emerge in West Africa by a policy-driven phase change in agroecology? in Neoliberal Ebola 1–12 (Springer, 2016).

200. Rulli, M. C., Santini, M., Hayman, D. T. & D’Odorico, P. The nexus between forest fragmentation in Africa and Ebola virus disease outbreaks. Sci. Rep. 7, (2017).

201. Carlson, C. J., Dougherty, E. R. & Getz, W. An ecological assessment of the pandemic threat of Zika virus. PLoS Negl Trop Dis 10, e0004968 (2016).

